# A DNA barcode library for *Culex* mosquitoes (Diptera: Culicidae) of South America with the description of two cryptic species of subgenus *Melanoconion*

**DOI:** 10.1101/2024.09.04.611342

**Authors:** Stanislas Talaga, Amandine Guidez, Benoît de Thoisy, Anne Lavergne, Romuald Carinci, Pascal Gaborit, Jean Issaly, Isabelle Dusfour, Jean-Bernard Duchemin

## Abstract

The genus *Culex* is one of the most diverse in the world and includes numerous known vector species of parasites and viruses to humans. Morphological identification of *Culex* species is notoriously difficult and rely mostly on the examination of properly dissected male genitalia which largely prevents female and immature identification during entomological, ecological or arboviral surveys. The aims of this study were (i) to establish a DNA barcode library for *Culex* mosquitoes of French Guiana based on the mitochondrial gene cytochrome c oxidase I (COI) marker, (ii) to compare three approaches of molecular delimitation of species to morphological identification, and (iii) to test the effectiveness of the COI marker at a broader geographical scale across South America. Mosquitoes used in this study were sampled in French Guiana between 2013 and 2023. We provide 246 COI sequences for 90 morphologically identified species of *Culex*, including five new country records and two newly described species. Overall, congruence between morphological identification and molecular delimitations using the COI barcode were high. The Barcode of Life Data clustering approach into Barcode Index Numbers gives the best result in terms of species delimitation, followed by the muti-rate Poisson Tree Processes and the Assemble Species by Automatic Partitioning methods. Inconsistencies between morphological identification and molecular delimitation can be explained by introgression, incomplete lineage sorting, imperfect taxonomy or the effect of the geographical scale of sampling. This increases by almost two-fold the number of mosquito species for which a DNA barcode is available in French Guiana, including 75% of the species of *Culex* currently known in the territory. Finally, this study confirms the usefulness of the COI barcode in identifying *Culex* mosquitoes of South America, but also points the limits of this marker for some groups of species within the subgenera *Culex* and *Melanoconion*.

## Introduction

The genus *Culex* is one of the most diverse in the world and includes numerous known vector species of parasites and viruses to humans. *Culex* mosquitoes are also of particular concern in view of the threat of emerging diseases in relation to global warming and environmental change [1]. In tropical America, this genus surpasses any other one in term of diversity and several arboviral studies pointed out its importance as vector of viruses. For example, a ten-years arboviral survey in Trinidad resulted in the isolation in *Culex* mosquitoes of 320 out of the 473 (68%) virus isolates [2]. In the Peruvian Amazon, species of *Culex* accounted for 57% of the mosquitoes collected, but 87% of the virus isolations were made from this genus [3]. In northeastern Amazonia, no fewer than fifteen arboviruses have been detected among *Culex* species of French Guiana [4]. To date, this oversea territory of France account for 113 nominal species of *Culex* classified among eight subgenera, representing more than 45% of the total number of recorded mosquito species [5; 6; 7; 8]. The most speciose subgenera of *Culex* were divided into informal infrasubgeneric groups that variously include Sections, Groups, Subgroups, Series and Complexes [1]. Morphological identification of *Culex* species is notoriously difficult and rely mostly on the examination of properly dissected male genitalia which largely prevents female and immature identification during entomological, ecological or arboviral surveys [9; 10; 11; 12].

Twenty years ago, Hebert et al. [13] established that the mitochondrial gene cytochrome c oxidase I (COI) can serve as the core of a global bioidentification system for animals. Since then, this taxon “barcode” was widely used successfully in diverse taxonomic groups (reviewed in [14)), including mosquitoes [15]. In South America, studies that have documented COI sequences for *Culex* mosquitoes are relatively scarce and gave mixed results as regard to delimitation and identification of species. One of the first study providing COI barcodes for South American *Culex* was part of an inventory of the mosquitoes of the Yasuni National Park of eastern Amazonian Ecuador [16]. Only five *Culex* species were included in this study but all of them were successfully delineated by this marker. In a series of articles dealing with the taxonomy of *Culex* (*Culex*) in South America, Laurito et al. [17; 18; 19] provided and analyzed COI sequences for 24 nominal species collected in Argentina and Brazil. Among this subgenus, the COI barcode barely identified 70% of the included species. In Colombia, sequences of 15 species/morphospecies of *Culex* from three ecosystems of the Andes permitted to delimit most taxa, except in the subgenus *Culex* [20]. Another important contribution was made available by Torres-Gutierrez et al. [21], which provided 120 COI sequences for 48 species/morphospecies of *Culex* (*Melanoconion*) from Brazil. These authors obtained coherent delimitation except in few cases where morphological species/morphospecies were split in more than one molecular unit [21]. Unfortunately, their dataset contains approximately 40% of female specimens for which the identification is inherently questionable. Recently, the barcode library for mosquitoes of Argentina was updated with additional species, including *Cx. amazonensis* (Lutz) and four species of *Culex* (*Melanoconion*) from the north-central part of the country [22]. All of them were correctly delimited by their DNA barcode and species of subgenus *Melanoconion* also clustered with specimens from Brazil.

A few years ago, we initiated a molecular database for barcoding and metabarcoding of mosquito species in French Guiana [23]. Overall, this study confirmed the effectiveness of both the COI and 16S markers in delimiting and identifying Guianese mosquitoes. Nevertheless, the genus *Culex* was largely overlooked as only 12% of the *Culex* species known in the territory were included. The aims of the present study were (i) to improve the taxonomic coverage of *Culex* in the barcode library of mosquitoes of French Guiana, (ii) to compare three approaches of molecular delimitation of species to morphological identification, and (iii) to test the effectiveness of the COI marker at a broader geographical scale across South America.

## Material and Methods

### Sampling and a priori identification

*Culex* mosquitoes were sampled using a wide range of methods and traps between 2013 and 2023 in French Guiana. A great diversity of natural habitats was sampled from the coastal plain to the upland terra firme forest, including inhabited and variously anthropized areas. Immature individuals were individually reared in the laboratory until emergence and associated larval and pupal exuviae were conserved in 70% alcohol whenever possible. In most cases, morphological identification has been done by microscopic observation of properly prepared, dissected, and mounted genitalia of males. After separating the last abdominal segments from the rest of the body, male genitalia were cleared in a 10% KOH solution for 2 hours at 40 °C, then stained in a 1% acid fuchsin solution for 5 minutes at room temperature, and finally dissected in a solution of Marc André [24]. Once mounted in Euparal between slide and coverslip, male genitalia were examined using an EVOS FL-Auto inverted microscope (Thermo Fisher Scientific Inc., Waltham, MA, USA). Identification was made using an extensive amount of taxonomic literature, including original description of species and taxonomic revisions like Bram [9], Duret [25], Valencia [10], Berlin and Belkin [11], Sirivanakarn [12], Sallum and Forattini [26], Pecor et al. [27], Sá et al. [28], and Sá et al. [29]. Specimens were selected to increase as much as possible the taxonomic and geographic coverage of the dataset.

### DNA extraction and sequencing

Total DNA of selected specimens was extracted from three legs or the abdomen (except male genitalia) of each adult, or the head of larval specimens. PCR amplification was performed using LCO1490 and HCO2198 primers [30], which are the standard for amplifying the 658-bp barcode region at the 5′ end of the COI gene [13]. The detailed protocol of amplification and sequencing that we used can be found in previous works [7; 23]. This article and its nomenclatural acts were registered in Zoobank (https://www.zoobank.org/). The life science identifier (LSID) of the article is: urn:lsid:zoobank.org:pub:C5D9E279-8B6E-45F3-86A1-8D6B78394715. Voucher specimens and DNA templates (including holotypes and paratypes of new species) are stored at the Institut Pasteur de la Guyane (IPG).

Forty-two specimens of *Culex* belonging to 14 morphologically identified species/morphospecies were already included in Talaga et al. [23]. These specimens were re-examined with regard to the knowledge acquired during a recent review of the *Culex* mosquitoes of French Guiana [6]. As a result, specimens MB1#0038, 0039, MB1#0225‒0227, and MB1#0810, 0811 turned out to be misidentifications of *Cx. urichii* (Coquillett), *Cx. nigripalpus* Theobald and *Cx. originator* Gordon & Evans, instead of *Cx. infoliatus* Bonne-Wepster & Bonne, *Cx. mollis* Dyar & Knab, and *Cx. imitator* Theobald, respectively. In addition, morphospecies named *Culex* sp.stI, sp.stJ, sp.stK and sp.stL have been respectively identified as the following nominal species: *Culex secundus* Bonne-Wepster & Bonne, *Cx. comminutor* Dyar, *Cx. imitator* and *Cx. putumayensis* Matheson. The taxonomic identification of these voucher specimens has been modified accordingly in BOLD and GenBank databases. Finally, five specimens initially identified as *Cx. imitator* (ST1#0310, 0311) and *Cx. stonei* Lane & Whitman (MB1#0173, 0241, 0242) were not included here because their identification could not be ascertained by any male genitalia.

### Molecular delimitation of species

Additional contig sequences were built with CodonCode before to be uploaded to the Barcode of Life Data Systems (BOLD) [31] as part of the FGMOS project, which gathers all the barcoding data available on the mosquitoes of French Guiana. BOLD accession numbers of specimens and barcode index numbers (BINs) are provided throughout the manuscript. A Maximum Likelihood (ML) phylogenetic analysis of the COI marker using the Kimura’s two-parameter model with defaults settings was conducted in Mega X [32]. A sequence of a specimen of *Chagasia bonneae* Root (WRBUE110-10) was used as outgroup, and nodal support was assessed using a bootstrap procedure under 1,000 replications. Afterward, morphological identification of species was compared with molecular delimitation using the BOLD BINs method and two standalone methods: the Assemble Species by Automatic Partitioning (ASAP) distance-based method [33] and the multi-rate Poison Tree Processes (mPTP) tree-based method [34]. ASAP divides species partitions based on pairwise genetic distances. ASAP also computes a probability of panmixia (p-val), a relative gap width metric (W), and ranked results by the ASAP score: the lower the score, the better the partitioning [33]. The PTP method is a phylogeny-aware approach that take the evolutionary relationships of the sequences into account. The multi-rate PTP (mPTP) model incorporate the potential divergence in intraspecific diversity and thereby it can better accommodate the sampling- and population-specific characteristics of a broader range of empirical datasets [34].

Finally, our dataset was compared to the others COI sequences of *Culex* available in South America using the BIN delimitation in BOLD. Sequences from Argentina (Laurito et al. [17; 22], Brazil [17; 21], Colombia [20] and Ecuador [16] were included from BOLD.

## Results

### Species identification and delimitation

A total of 246 specimens of *Culex* mosquitoes belonging to 8 subgenera and 90 morphologically identified species from 44 sampling sites were included in the analyses (Figure 1, S1 Table, Supporting information). Readers interested in the authorship of the species sequenced in this study should consult S1 Table. Selected specimens were mostly represented by males with dissected genitalia (78%), followed by larvae (14%) and females (8%). Among them, 80 species were represented by two or more specimens (up to five), but ten species were only represented by one specimen. The dataset included five new country records and two newly described species. The new country records included, *Cx. bibulus* (collected in the Amerindian villages of Camopi and Twenké, and Savanes de Passoura), *Cx. galindoi* (collected along a small river inside the Réserve Naturelle Nationale de La Trinité), *Cx. johnnyi* (collected along the Crique Gabaret, a medium-sized river tributary of the Oyapock), *Cx. longistriatus* (collected along a large river under tidal influence at Roura) and *Cx. ocossa* (collected close by a coastal marsh at Pointe Macouria).

**Fig 1.**
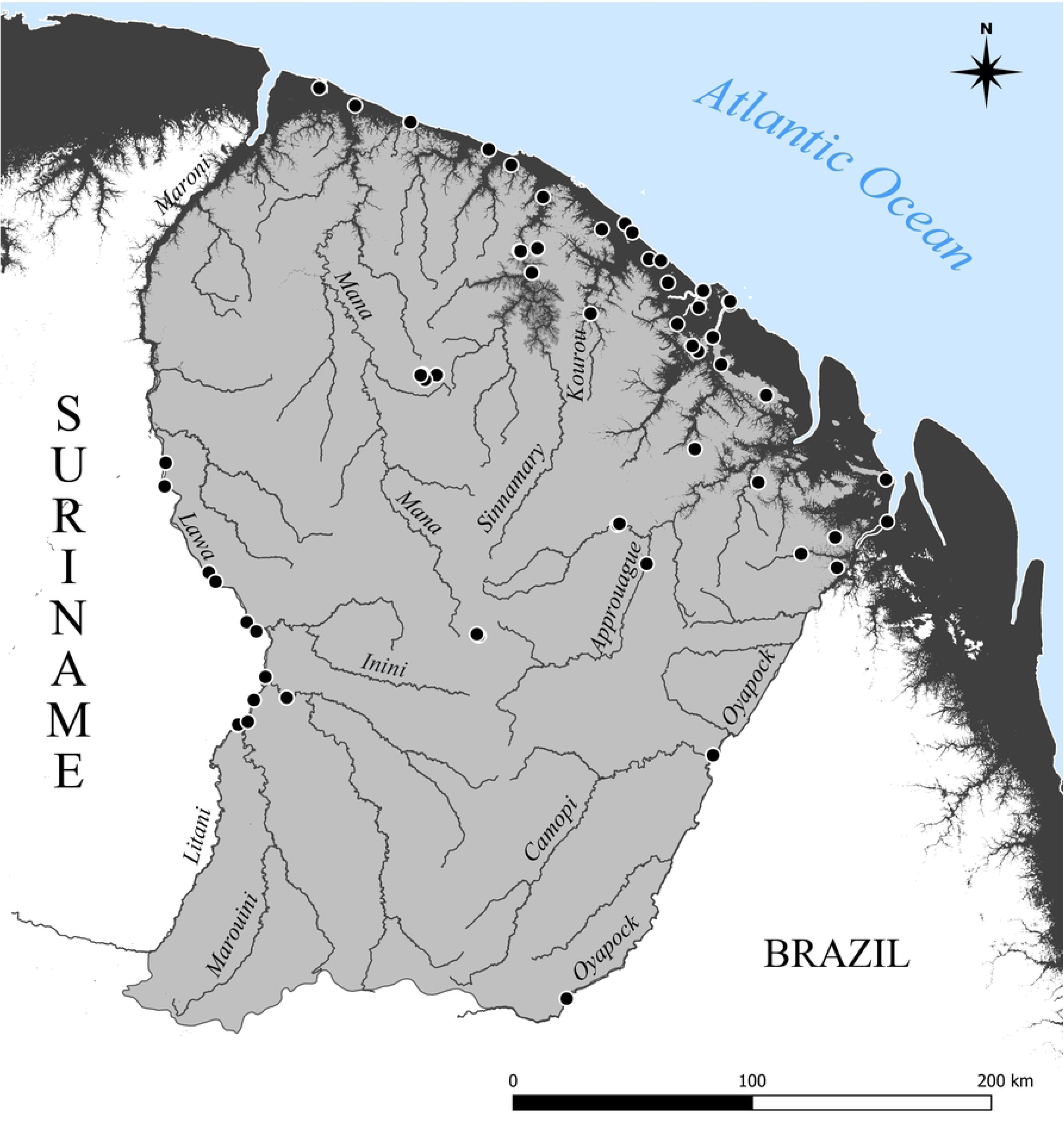
Map showing the distribution of sampling localities of the *Culex* (Diptera: Culicidae) specimens from which the COI barcode was sequenced in this study. French Guiana is coarsely divided into the coastal plain composed of a mosaic of mangroves, marshes, swamps, savannas and forests (dark gray; below 30 m a.s.l.) and upland terra firme forest (light gray; above 30 m a.s.l.). The main rivers are indicated.

The BOLD clustering approach allowed to distinguish 87 BINs out of the 90 morphologically identified species (Figure 2A‒C, S2 Table, Supporting information). Among them, 44 BINs were new to BOLD, the 43 remaining BINs included sequences already present in BOLD, including five species of *Culex* recently described from French Guiana [7; 8]. The results of the clustering approach into BINs were largely congruent with the morphological identification. However, 17% of the species were grouped into BINs with mixed nominal species, preventing their accurate identification. In details, we found five cases where two or more nominal species (up to four) were clustered into a single BIN; namely: BIN ADK0770 clustered sequences of *Cx. batesi* (N=1) and *Cx. evansae* (N=2), BIN AEE2103 clustered sequences of *Cx. contei* (N=3), *Cx. phlogistus* (N=3), and *Cx. serratimarge* (N=3), BIN AAN3636 clustered sequences of *Cx. brevispinosus* (N=2), *Cx. surinamensis* (N=2), and *Cx. usquatus* (N=7), BIN AAF1735 clustered sequences of *Cx. declarator* (N=3), *Cx. mollis* (N=2), and *Cx. nigripalpus* (N=5), and BIN AEE6759 clustered sequences of *Cx. creole* (N=4), *Cx. eastor* (N=4), *Cx. hutchingsae* (N=3), and *Cx. idottus* (N=3). Moreover, in seven cases, nominal species were split into two BINs; namely: *Culex bastagarius* (BINs AEE3102 and AFI9514), *Cx. foliafer* (BINs AEE1181 and AEE1182), *Cx. inadmirabilis* (BINs AFJ0561 and AFJ0562), *Cx. bibulus* (BINs AEE1543 and AFI9809), *Cx. organaboensis* (BINs AFJ0416 and AFT5569), *Cx. rabelloi* (BINs ADE6009 and AEW5154), and *Cx. theobaldi* (BINs ADK5539 and AEE9553).

**Fig 2.**
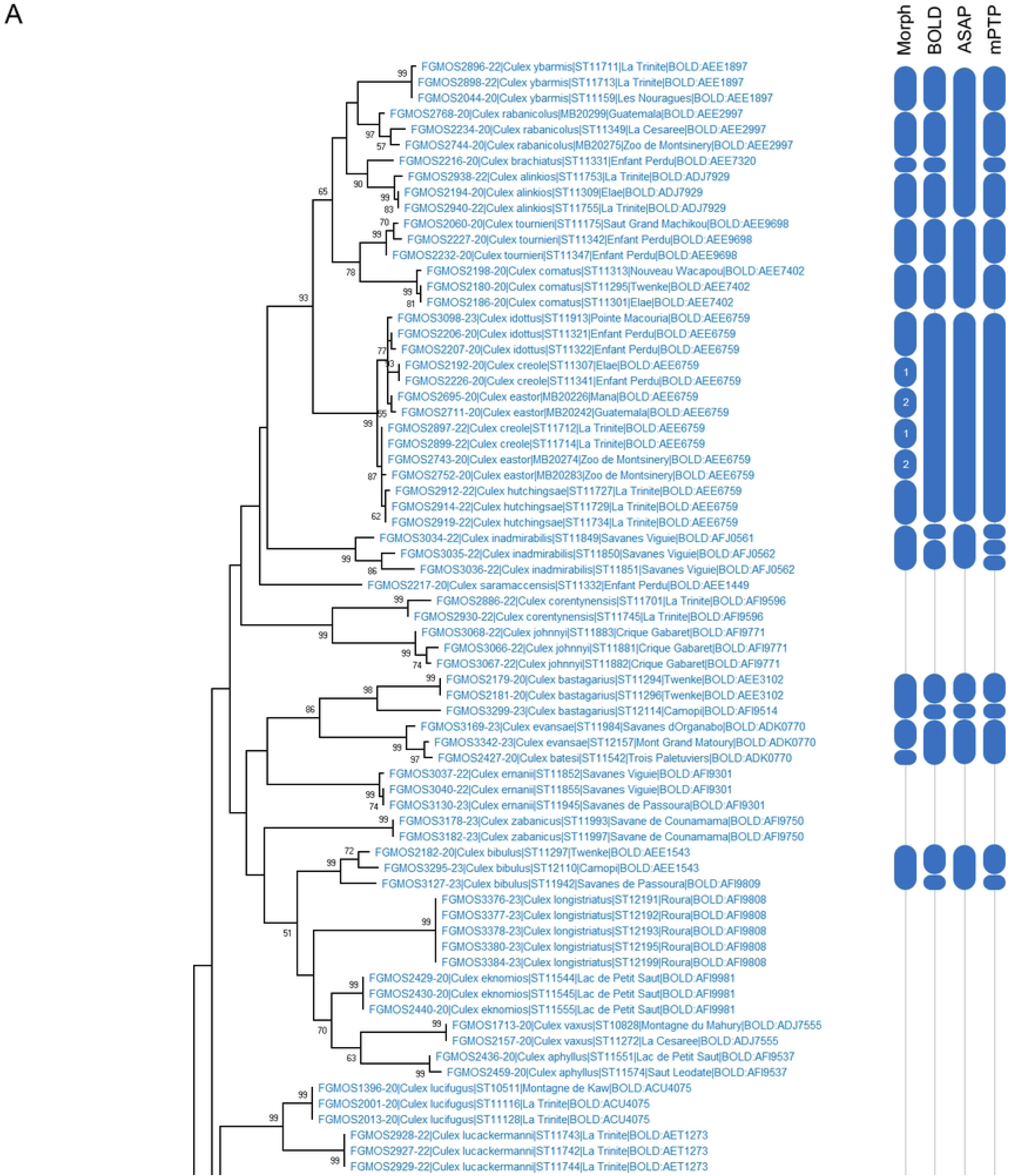
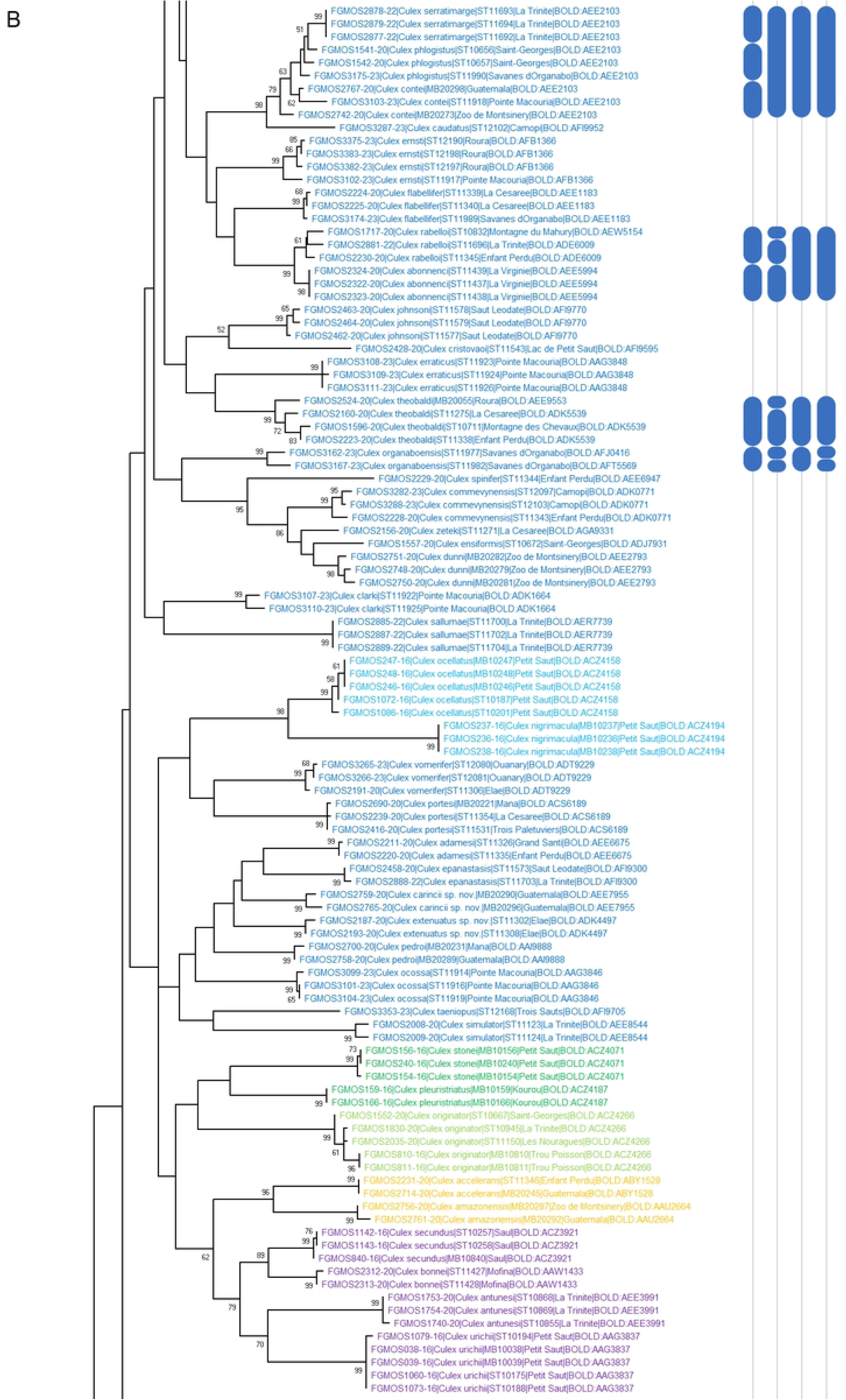
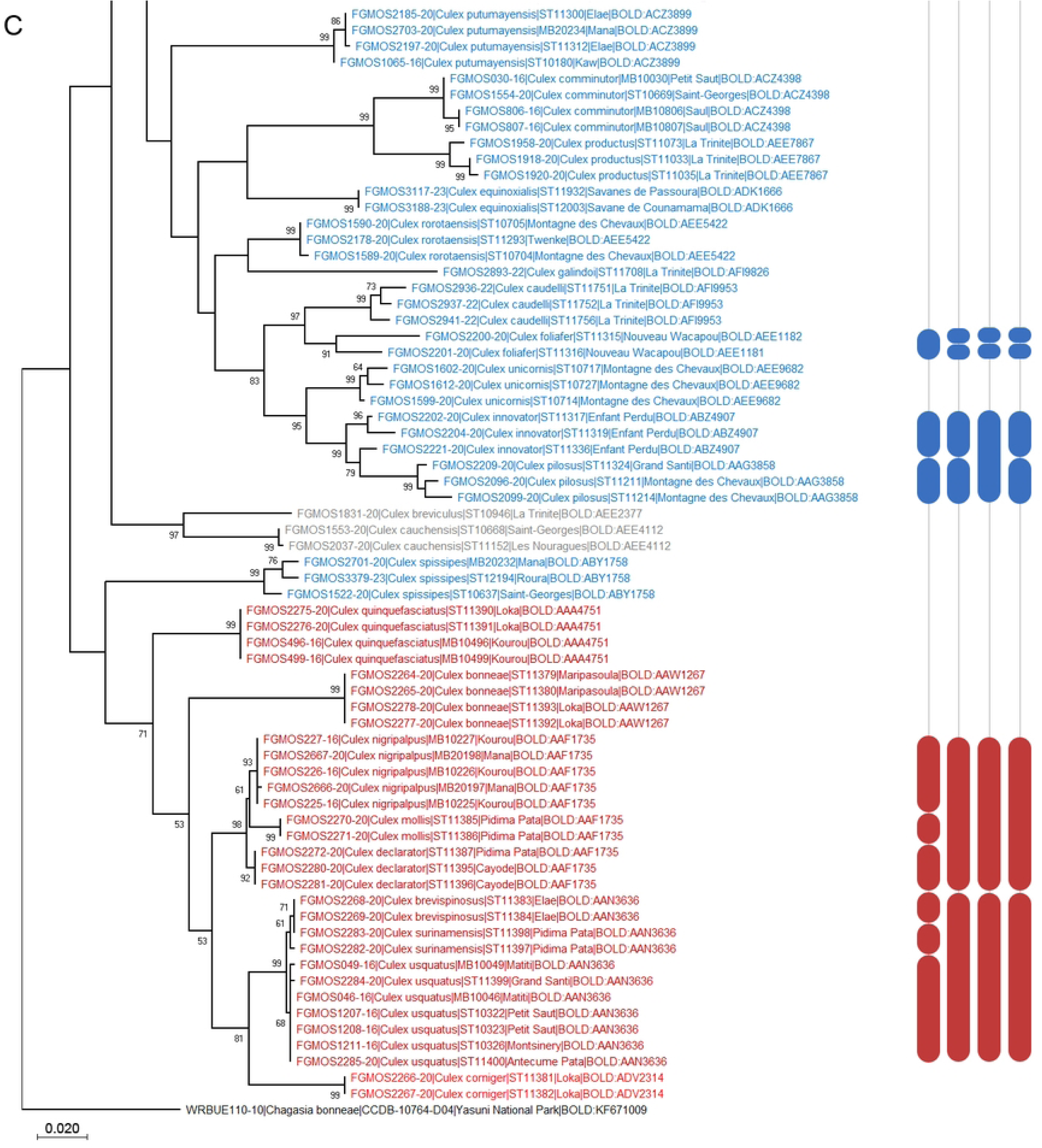
Maximum Likelihood (ML) phylogenetic analysis of the COI dataset of *Culex* mosquitoes (Diptera: Culicidae) from French Guiana (A‒C). For each specimen, we indicated the BOLD specimen code, the morphological identification, the original specimen code, the sampling locality, and the BOLD Barcode Index Number (BIN). Inconsistencies between morphological identification and molecular delimitation using the BOLD BIN, the Assemble Species by Automatic Partitioning (ASAP), and the multi-rate Poison Tree Processes (mPTP) methods are figured in column on the right side. *Culex* specimens are color coded by subgenus as follows: *Aedinus* in orange, *Anoedioporpa* in light green, *Carrollia* in purple, *Culex* in dark red, *Melanoconion* in blue, *Microculex* in green, *Phenacomyia* in red, and *Tinolestes* in grey. *Culex* formerly placed in the Ocellatus Section, presently without subgeneric placement, are colored in light blue. Numbers indicate split specimens belonging to the same species. Bootstrap support values above 50% are indicated near the nodes and a sequence of *Chagasia bonneae* Root was used as outgroup.

The ASAP method was mostly congruent with the results of the BOLD delimitation into BINs. However, the best model retrieved only 76 partitions among the COI dataset with 28% of the species included into partitions with mixed nominal species. Compared to the BIN delimitation, we find four more cases where two or more nominal species (up to four) were grouped in the same partition; namely: *Culex abonnenci* (N=3) and *Cx. rabelloi* (N=3) were grouped in the same subset, *Cx. alinkios* (N=3), *Cx. brachiatus* (N=1), *Cx. rabanicolus* (N=3) and *Cx. ybarmis* (N=3) were grouped in the same subset, *Cx. comatus* (N=3) and *Cx. tournieri* (N=3) were grouped in the same subset, and *Cx. innovator* (N=3) and *Cx. pilosus* (N=3) were grouped in the same subset. On the other hand, this method retrieved *Cx. bibulus*, *Cx. inadmirabilis*, *Cx. organaboensis*, and *Cx. theobaldi* in molecular subsets concordant with their morphological identification, meaning one partition instead of several BINs per species.

The mPTP method was highly congruent with the results of the BOLD delimitation into BINs and retrieved the same number of molecularly delimited species (87). However, 19% of the species were grouped into subsets with mixed nominal species. Compared to the BIN delimitation, we find one more case where two species were grouped in the same subset; namely: *Culex abonnenci* (N=3) and *Cx. rabelloi* (N=3) were grouped in the same molecular unit, as with the ASAP method. In three other cases BINs were split into two or three molecular units; namely: *Culex corentynensis* (BIN AFI9596) was split into two molecular units, *Cx. inadmirabilis* (BINs AFJ0561 and AFJ0562) was split into three molecular units, and *Cx. innovator* (BIN ABZ4907) was split into two molecular units. On the other hand, specimens of *Cx. theobaldi* (BINs AEE9553 and ADK5539) were grouped in the same molecular unit.

### Species description

Two cryptic species turned out to be new after COI sequencing and in-depth investigations, including examination of primary type specimens of morphologically close species. In order to help future works, they are formally described and named below.

#### *Culex* (*Melanoconion*) *carincii* Talaga & Duchemin, sp. nov

LSID: urn:lsid:zoobank.org:act:BEA55CAB-4766-408A-9F7B-788CD6768B99 BIN: AAE7955

**Male.** Habitus briefly examined. Hindtarsomeres 1‒5 completely dark. *Genitalia* (Figures 3A‒G): Tergum VIII with a deep V-shaped emargination separating the 2 lateral lobes. Tergum IX with 2 distinct lobes, conical to mound-like in shape, widely separated, bearing 6‒8 setae. Gonocoxite conical; inner margin moderately concave; ventrolateral setae strongly developed; lateral surface with 14‒19 small, scattered setae (lsp) at level of subapical lobe; proximal part of ventrolateral surface with scales. Subapical lobe clearly divided into 2 divisions. Proximal division of subapical lobe with an apical infundibular and hyaline expansion, 2 robust, sinuous, apically hooked setae (setae *a* and *b*) at apex, a subapical hyaline, broad, hooked-falciform seta and 4‒6 strong, pointed setae from base to level of insertion of the hooked-falciform seta; a patch of 9‒12 short setae inserted mesally at base of distal surface. Distal division of subapical lobe divided into 2 divergent arms, the proximal arm more robust than the distal arm bearing 3 apical setae, 1 long hooked seta (*h*), 1 short saber-like seta (*s*), and 1 long foliform seta (*l*) inserted clearly proximal to setae *h* and *s*; distal arm bearing 5 setae gradually inserted from apex to base, 1 long, curved, saber-like seta (*s*), and 4 appressed flattened setae (*f*), the most basal conspicuously shorter. Gonostylus slender, curved and slightly constricted at midlength, subapical portion barely enlarged on lateral view; 1 long seta inserted at proximal 0.25 on 1 out of the 4 gonostylus of the type series; ventral surface with conspicuous crest extending from apical snout to enlarged subapical portion; apical snout tapered to a truncate apex; gonostylar claw short, leaflike, highly broadened apically; ventral margin with 2 setae before gonostylar claw, distal seta conspicuously longer than proximal seta. Phallosome with lateral plates and aedeagal sclerites equivalent in length; aedeagal sclerite broad, curved in lateral view and broadly connected to base of lateral plate; distal part of lateral plate without median process, sternal and tergal processes present; apical sternal process short, laterally curved, pointed at apex; apical tergal process longer than apical sternal process, pointed and directed dorsolaterally. Proctiger elongate; paraproct narrowed distally, expanded basally, crown a row of about 7, 8 short simple blades. Cercal sclerite long and narrow with 2 cercal setae. Basal plate, paramere and tergum X as figured.

**Fig 3.**
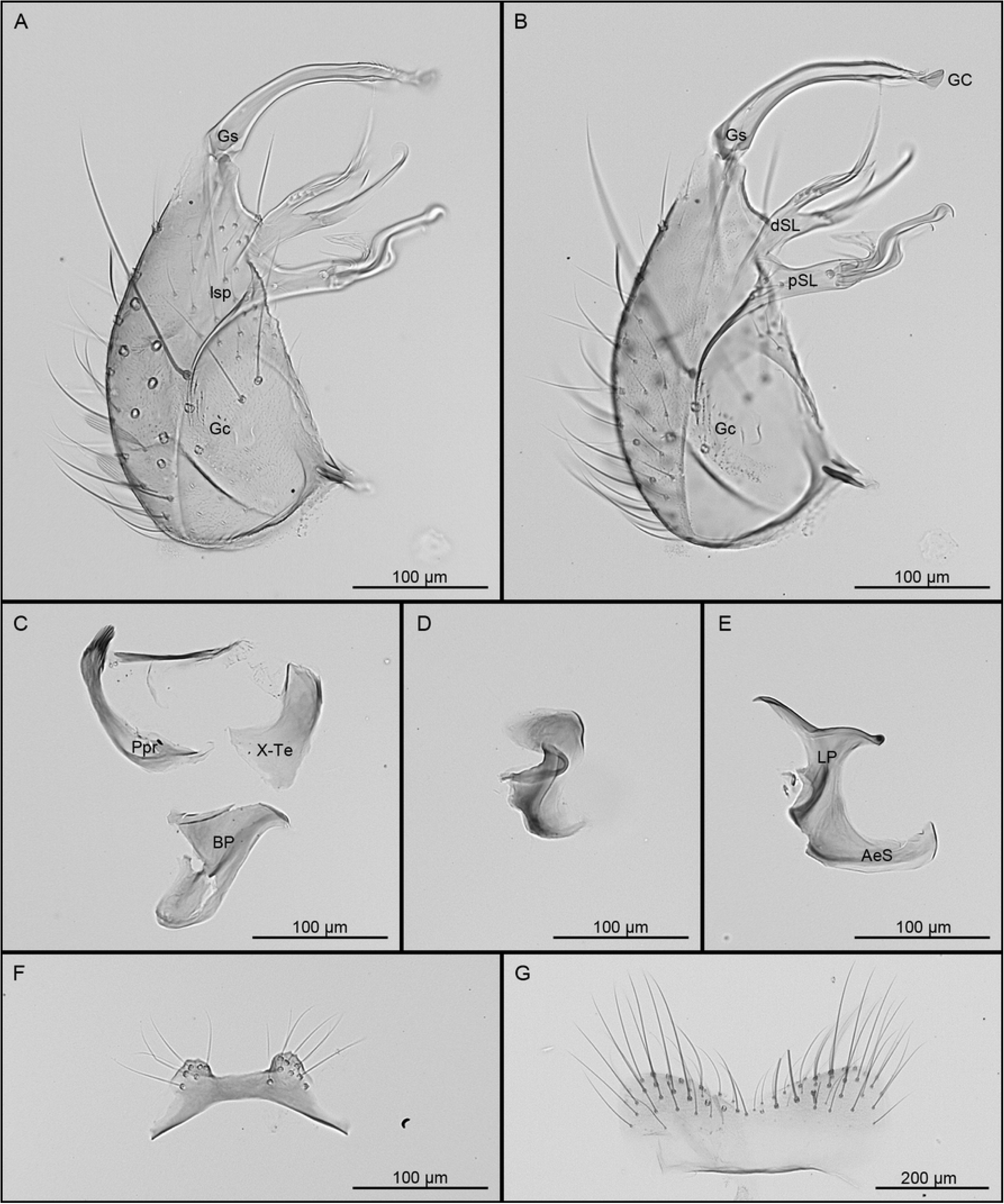
**Male genitalia of *Culex* (*Melanoconion*) *carincii* sp. nov.** A, Gonocoxopodite, lateral aspect; B, gonocoxopodite, mesal aspect; C, paraproct, tergum X and basal plate, in lateral views; D, paramere, lateral view; E, lateral plate and aedeagal sclerite, lateral views; F, tergum IX; G, tergum VIII. Morphological structures are abbreviated as follow: AeS, aedeagal sclerite; BP, basal plate; dSL, distal division of subapical lobe; Gc, gonocoxite; GC, gonostylar claw; Gs, gonostylus; LP, lateral plate; lsp, lateral setal patch; Ppr, paraproct; pSL, proximal division of subapical lobe; X-Te, tergum X.

**Etymology.** This species is dedicated to our dear colleague Romuald Carinci for his unflagging enthusiasm working on mosquitoes at the Institut Pasteur de la Guyane.

**Bionomics.** Nothing is known about the bionomics of *Cx. carincii*. Adult males were collected using CDC light traps (supplemented with black light) placed at 1.5 m above ground and operated from 1700 to 0700 h in a lowland forest patch surrounded by swamps and marshes on the coastal plain of French Guiana.

**Distribution.** *Culex carincii* is only know from the type locality.

**Type material.** *Holotype*: Adult male in 96% ethanol with dissected genitalia mounted on a microscope slide (specimen number MB2#0290, BOLD: FGMOS2759-20, GenBank: [waiting for accession creation]), FRENCH GUIANA: Guatemala (52.626420° W, 5.136090° N, 3 m above sea level), 22-VI-2017, S. Talaga and R. Carinci, IPG. *Paratypes*: Two adult males in 96% ethanol with dissected genitalia mounted on separate microscope slides (specimen numbers MB2#0296, BOLD: FGMOS2765-20, GenBank: [waiting for accession creation], MB2#0291, BOLD: FGMOS2760-20), same collection data as the holotype, IPG.

**Other material examined.** Holotype male genitalia of *Cx. crybda* Dyar (specimen number USNMENT01935104), USNM.

#### *Culex* (*Melanoconion*) *extenuatus* Talaga & Duchemin, sp. nov

LSID: urn:lsid:zoobank.org:act:D0E621BA-3C9A-4928-8965-4C0A63A8C42D BIN: ADK4497

**Male.** Habitus briefly examined. Hindtarsomeres 1‒4 with conspicuous white rings on base and apex, 5 white. *Genitalia* (Figures 4A‒G): Tergum VIII with a deep V-shaped emargination separating the 2 lateral lobes. Tergum IX with 2 distinct lobes, conical to mound-like in shape, widely separated, bearing 8‒14 setae. Gonocoxite conical; inner margin moderately concave; ventrolateral setae strongly developed; lateral surface with 13‒17 small, scattered setae (lsp) at level of subapical lobe; proximal part of ventrolateral surface with scales. Subapical lobe clearly divided into 2 divisions. Proximal division of subapical lobe with an apical infundibular and hyaline expansion, 2 robust, sinuous, apically hooked setae (setae *a* and *b*) at apex, a subapical hyaline, broad, hooked-falciform seta and 5, 6 strong, pointed setae from base to level of insertion of the hooked-falciform seta; a patch of 10‒14 short setae inserted mesally at base of distal surface. Distal division of subapical lobe divided into 2 divergent arms, the proximal arm bearing 3 apical setae, 1 long hooked seta (*h*), 1 shorter saber-like seta (*s*), and 1 long foliform seta (*l*) inserted proximal to setae *h* and *s*; distal arm bearing 5 apical setae, 1 long, curved, saber-like seta (*s*), and 4 appressed flattened setae (*f*). Gonostylus slender, curved at midlength, subapical portion enlarged on lateral view; ventral surface with inconspicuous crest extending from apical snout to enlarged subapical portion; apical snout tapered to a truncate apex; gonostylar claw short, leaflike, broadened apically; ventral margin with 2 setae before gonostylar claw, distal seta conspicuously longer than proximal seta. Phallosome with lateral plates and aedeagal sclerites equivalent in length; aedeagal sclerite broad, curved in lateral view and broadly connected to base of lateral plate; distal part of lateral plate without median process, sternal and tergal processes present; apical sternal process short, laterally curved, pointed at apex; apical tergal process longer than apical sternal process, pointed and directed dorsolaterally. Proctiger elongate; paraproct narrowed distally, expanded basally, crown a row of about 7, 8 short simple blades. Cercal sclerite long and narrow with 2, 3 cercal setae. Basal plate, paramere and tergum X as figured.

**Fig 4.**
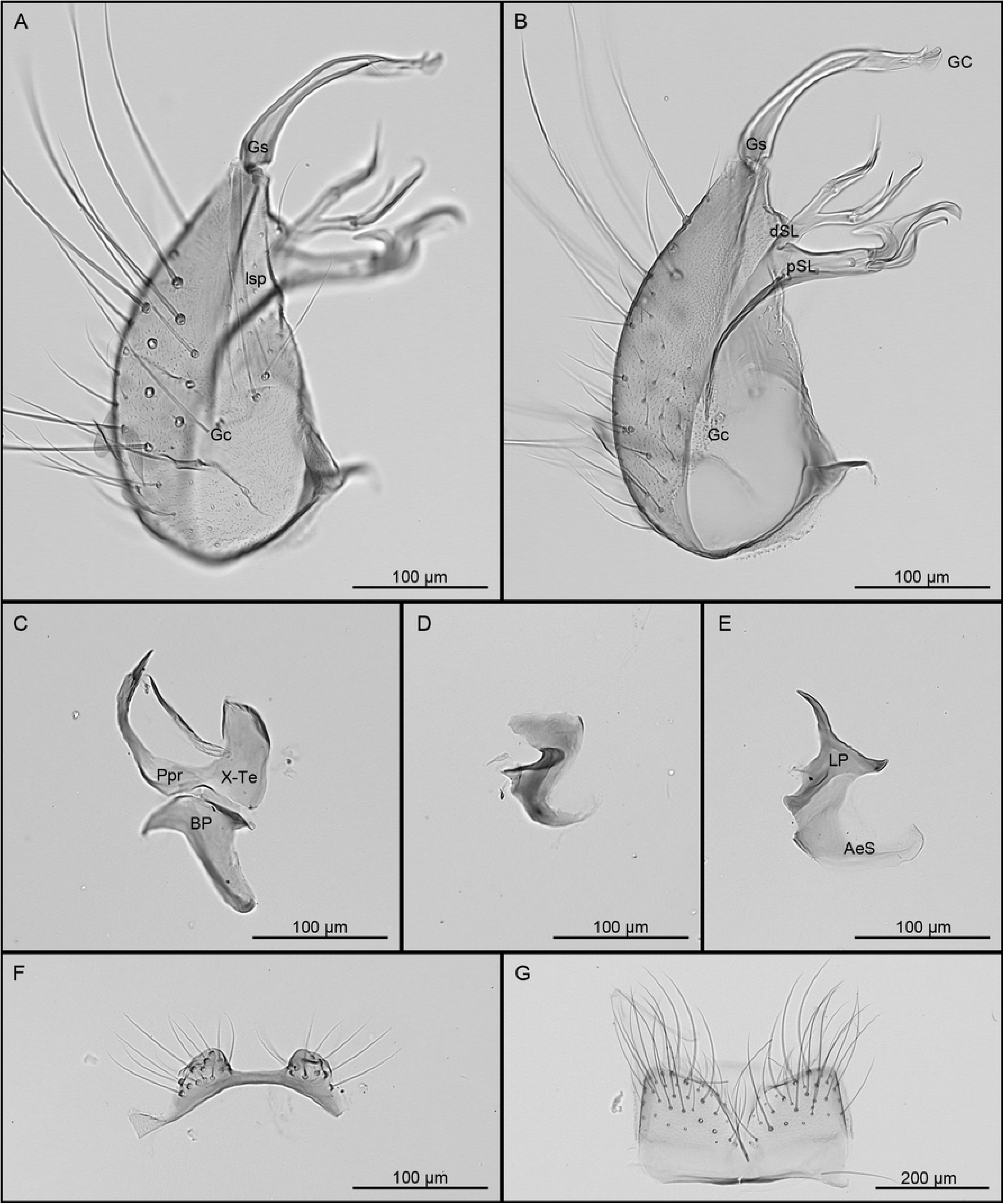
**Male genitalia of *Culex* (*Melanoconion*) *extenuatus* sp. nov.** A, Gonocoxopodite, lateral aspect; B, gonocoxopodite, mesal aspect; C, paraproct, tergum X and basal plate, in lateral views; D, paramere, lateral view; E, lateral plate and aedeagal sclerite, lateral views; F, tergum IX; G, tergum VIII. Morphological structures are abbreviated as follow: AeS, aedeagal sclerite; BP, basal plate; dSL, distal division of subapical lobe; Gc, gonocoxite; GC, gonostylar claw; Gs, gonostylus; LP, lateral plate; lsp, lateral setal patch; Ppr, paraproct; pSL, proximal division of subapical lobe; X-Te, tergum X.

**Etymology.** *Culex extenuatus* is named for the thinned arms (proximal and distal) of the distal division of the subapical lobe of the gonocoxite relatively to the other species of the Pedroi Subgroup of the Spissipes Section of *Culex* subgenus *Melanoconion*.

**Bionomics.** Nothing is known about the bionomics of *Cx. extenuatus*. Adult males were collected using CDC light traps (supplemented with black light or not) placed at 1.5 m above ground and operated from 1700 to 0700 h in the secondary vegetation surrounding the Amerindian village of Elaé on the Lawa River.

**Distribution.** *Culex extenuatus* is know from the type locality and few inland localities in French Guiana, namely: Cacao, Crique Gabaret, and Grand Santi.

**Type material.** *Holotype*: Adult male in 96% ethanol with dissected genitalia mounted on a microscope slide (specimen number ST1#1302, BOLD: FGMOS2187-20, GenBank: [waiting for accession creation]), FRENCH GUIANA: Lawa River, Elaé (54.048683° W, 3.380118° N, 100 m above sea level), 22-III-2018, S. Talaga, IPG. *Paratype*: One adult male in 96% ethanol with dissected genitalia mounted on a microscope slide (specimen number ST1#1308, BOLD: FGMOS2193-20, GenBank: [waiting for accession creation]), same collection data as the holotype, IPG.

**Other material examined.** Three adult males in 96% ethanol with dissected genitalia mounted on separate microscope slides, FRENCH GUIANA: Grand Santi (54.381119° W, 4.270310° N, 50 m above sea level), 28-III-2018, S. Talaga, IPG (specimen number ST1#1325); Cacao (52.466964° W, 4.582039° N, 3 m above sea level), 23-VII- 2019, O. Romoli and K. Heu, IPG (specimen number ST1#1567); Crique Gabaret (51.915540° W, 3.925270° N, 25 m above sea level), 19-II-2021, S. Talaga, IPG (specimen number ST1#1873). Holotype male genitalia of *Cx. pedroi* Sirivanakarn & Belkin (specimen number USNMENT01935483), USNM.

### COI sequences of *Culex* across South America

Comparison of our dataset with the others COI sequences of *Culex* available in South America (529 sequences of 150 species/morphospecies) revealed accurate matches, close matches and mismatches between nominal species and molecular BINs (S3 Figure, Supporting information). Comparison involving species absent of our dataset and morphospecies were not detailed in order not to lengthen the article too much.

In 18 cases, *Culex* species identified outside French Guiana clustered in the same BIN as our specimens of the same species. For the subgenus *Aedinus*, specimens of *Cx. amazonensis* from Argentina clustered in the same BIN (AAU2664) as our specimens from French Guiana. In *Carrollia*, specimens of *Cx. bonnei* and *Cx. urichii* from Ecuador clustered in the same BINs (AAW1433 and AAG3937, respectively) as our specimens from French Guiana. For the subgenus *Culex*, specimens of *Cx. quinquefasciatus* from Argentina clustered in the same BIN (AAA4751) as our specimens from French Guiana. Specimens of *Cx. declarator* and *Cx. nigripalpus* from Brazil and Colombia, and *Cx. mollis* from Argentina and Brazil clustered in the same BIN (AAF1735) as our specimens from French Guiana. Specimens of *Cx. surinamensis* from Brazil and a specimen of *Cx. usquatus* from Argentina clustered in the same BIN (AAN3636) as our specimens from French Guiana. For the subgenus *Melanoconion*, a specimen of *Cx. evansae* and a specimen of *Cx. ensiformis* from Brazil clustered in the same BINs (ADK0770 and ADJ7931, respectively) as our specimens from French Guiana. Specimens of *Cx. erraticus* and *Cx. lucifugus* from Colombia clustered in the same BINs (AAG3848 and ACU4075, respectively) as our specimens from French Guiana. Specimens of *Cx. clarki*, *Cx. commevynensis*, *Cx. equinoxialis* and a specimen of *Cx. vaxus* from Brazil clustered in the same BINs (ADK1664, ADK0771, ADK1666, and ADJ7555, respectively) as our specimens from French Guiana. Finally, specimens of *Cx. theobaldi* from Brazil clustered in the same BIN (ADK5539) as a most of our specimens from French Guiana.

In 14 cases we found mismatches where the same nominal species clustered in different BINs. At one exception, all these cases involved species of the subgenus *Melanoconion*. Among the Melanoconion Section, Specimens of *Cx. dunni* from Argentina and Brazil and *Cx. zeteki* from Brazil clustered in different BINs (ADJ7556 and AAZ5313, respectively) from those originating in French Guiana (AEE2793 and AGA9331, respectively). Specimens of *Cx. corentynensis* from Brazil clustered in a distant BIN (ADJ8504) from those originating in French Guiana (AFI9596). Specimens of *Cx. putumayensis* from Brazil clustered in a different but related BIN (ADJ8613) from those originating in French Guiana (ACZ3899). Specimens of *Cx. vaxus* from Argentina and some from Brazil clustered in different BINs (AEB4367, ADM0315, and ADK6634, respectively) than those originating in French Guiana (ADJ7555). Specimens of *Cx. aureonotatus* Duret & Barreto from Brazil clustered in the same BIN (AAG3848) as specimens of *Cx. erraticus* from French Guiana. Specimens of *Cx. serratimarge* from Brazil and Ecuador clustered in different BINs (ADK0466 and AAW1266, respectively) from those originating in French Guiana (AAE2103). Specimens of *Cx. idottus* from Argentina and Brazil and *Cx. eastor* from Brazil clustered in different BINs (ADG8243 and ADJ7929, respectively) from those originating in French Guiana (AEE6759). Specimens of *Cx. pilosus* from Brazil clustered in different BINs (ADJ8438 and ADT4465) than those originating in French Guiana (AAG3858). In the Spissipes Section, a specimen of *Cx. portesi* and specimens of *Cx. spissipes* and *Cx. vomerifer* from Brazil clustered in different BINs (AAZ3500, ADK0011 and ADK2041, respectively) to our specimens from French Guiana (ACS6189, ABY1758 and ADT9229, respectively). In the subgenus *Phenacomyia*, specimens of *Cx. corniger* from Colombia clustered in a different BIN (ABU8489) than those from French Guiana (ADV2314).

Finally, three specimens from other datasets clustered with sequences of different subgenera than their own. Two specimens of *Cx. conspirator* Dyar & Knab from Colombia (GenBank KM593054 and KM593048) clustered with species the subgenus *Culex*, and one specimen of *Cx. rabelloi* from Brazil (GenBank KX779859) clustered with sequences of *Cx. amazonensis* from Argentina and French Guiana. These specimens should be reviewed because their identifications are most likely erroneous.

## Discussion

Overall, congruence was high between morphological identification and molecular delimitations for *Culex* of French Guiana using the standard COI marker. The BOLD clustering approach into BINs gives the best result in terms of species delimitation, followed by the mPTP method and the ASAP method. In five cases, the three delimitation approaches grouped more than one nominal species in the same molecular unit: two cases included *Culex* (*Culex*) species (BINs AAF1735 and AAN3636), and three cases included *Culex* (*Melanoconion*) species (BINs ADK0770, AEE2103 and AEE6759), for a total of 15 species involved. Although these species are morphologically well-defined, this indicates that the COI marker does not contain enough information to accurately identify these species above their BIN. Such limit was already pointed out on a set of 22 *Culex* (*Culex*) species from Argentina and Brazil from which the COI barcode barely identified 70% of them [17]. Our results show that it can also apply in a lesser extent to the subgenus *Melanoconion* where the COI barcode allowed to accurately identify only 87% of them. On the other hand, some species hardly identifiable among the subgenus *Melanoconion*, even with properly dissected male genitalia, are unambiguously delimited based on the COI barcode. This is particularly true for morphologically close species of the Educator Group of the Melanoconion Section (for example, *Cx. bibulus*, *Cx. longistriatus* and *Cx. vaxus*) or of the Pedroi Subgroup of the Spissipes Section (for example, *Cx. crybda*, *Cx. extenuatus* and *Cx. pedroi*). *Culex bastagarius* and *Cx. foliafer* were the only two species as being split into more than one molecular unit by the three delimitation methods. This result can be a signal of close-related species, but careful examination did not permit to detect any morphological differences. Until more material become available to study, we interpret this delimitation as an artefact linked to sampling geographical scale or proof of separation then further introgression with persistence of mitochondrial DNA signal.

Most subgenera and some informal groups within the *Culex* genus are retrieved in well-supported monophyletic clades under the ML phylogenetic analysis of the COI barcode. This is the cases of subgenera *Aedinus* (two species included, 96% bootstrap value), *Anoedioporpa* (only one species included), *Carrollia* (four species included, 79% bootstrap value), *Tinolestes* (two species included, 97% bootstrap value), as well as the former Ocellatus Section presently without subgeneric placement represented here by *Cx. nigrimacula* and *Cx. ocellatus* (98% bootstrap value). Monophyly of the subgenera *Culex*, *Melanoconion* and *Microculex* are not supported statistically by our analysis. Species of *Culex* (*Culex*) included in the analysis clustered together in a clade supported at 71% of bootstrap value but this clade also included *Cx. corniger*, a member of the subgenus *Phenacomyia* Harbach and Peyton [35].

Overall, species of the Melanoconion and Spissipes sections of *Melanoconion* clustered together but their basal nodes were not supported by bootstrap values. Nevertheless, some informal groups within the Melanoconion Section were well- supported. This is the case of the Atratus Group (95%), the Conspirator Group (99%), and the Pilosus Group (83%), comprising the Caudelli Subgroup (97%) and the Pilosus Subgroup (95%). The monophyly of the Atratus and Pilosus groups were already shown by Torres-Gutierrez et al. [21; 36] using the COI barcode and two nuclear markers. The monophyly of the Educator Group as currently interpreted [29] is not sustained by our phylogenetic analysis of the COI marker. Only five out of the eight species included in this study (namely, *Cx. aphyllus*, *Cx. bibulus*, *Cx. eknomios*, *Cx. longistriatus*, and *Cx. vaxus*) were retrieved as monophyletic (51% bootstrap value). However, *Cx. cristovaoi*, *Cx. inadmirabilis* and *Cx. theobaldi* clustered far from this clade which suggests that they not belong to the Educator Group. Some other results are worth noting. A group of ten species originally classified among the Bastagarius Group (namely, *Cx. alinkios*, *Cx. brachiatus*, *Cx. comatus*, *Cx. creole*, *Cx. hutchingsae*, and *Cx. tournieri*) and the Intrincatus Group (namely, *Cx. eastor*, *Cx. idottus*, *Cx. rabanicolus*, and *Cx. ybarmis*) clustered together in a well- supported monophyletic clade (93% bootstrap value). In the male geniatalia, both groups are separated based on the shape of the apical median process of the lateral plate of the phallosome [12]. However, this morphological structure could show a complete intergradation from spinelike to broad quadrate, rendering it ineffective in separating species with intermediate shapes of apical median process. Three pairs of species originally placed in different informal groups were retrieved clustering together in statistically supported clades. This is the case of *Cx. corentynensis* clustering with *Cx. johnnyi* (99% bootstrap value). These species were originally placed in the Iolambdis Subgroup of the Bastagarius Group and Evansae Group, respectively [12]. However, male genitalia of both species show striking morphological similarities, especially regarding the setal arrangement of the distal subapical lobe of the gonocoxite [27]. Similarly, *Cx. cristovaoi* and *Cx. johnsoni* originally placed in the Educator Group and Intrincatus Group, respectively [12] clustered together (52% bootstrap value). The overall morphological similarity of these two species was noted in a recent revision of the Educator Group [29] and supported here by our analysis. Finally, *Cx. bastagarius* of the Bastagarius Subgroup of the Bastagarius Group clustered with two species of the Evansae Group (*Cx. evansae* and *Cx. batesi*) in a well-supported clade (86% bootstrap value). A similar result was found as regard to *Cx. bastagarius* and *Cx. evansae* collected in Brazil [21; 36]. All these results confirm the importance of genital structures in delimitation of *Culex* species but put into perspective the overriding importance of the lateral plate in the infrasubgeneric classification of the Melanoconion Section of the subgenus *Melanoconion*.

*Culex carincii* and *Cx. extenuatus* described herein belong to the Spissipes Section within the infrasubgeneric classification of the subgenus, based on the broad aedeagal sclerite, curved in lateral view and broadly connected to the base of the lateral plate [12; 26]. Both species belong to the Pedroi Subgroup of the Crybda Group, together with *Cx. adamesi*, *Cx. crybda*, *Cx. epanastasis*, *Cx. pedroi*, and *Cx. ribeirensis*. *Culex carincii* and *Cx. extenuatus* share with them all the distinguishing features of the Pedroi Subgroup [26]. This includes 1) the presence of scales on the proximal part of the ventrolateral surface of the gonocoxite, 2) the lateral plate of the phallosome without an apical median process, apical sternal and tergal processes present; the apical sternal process short, pointed, laterally curved; the apical tergal process long, nearly pointed, dorsolaterally directed; 3) the proximal division of the subapical lobe of the gonocoxite with an apical infundibular and hyaline expansion, a subapical broad hooked-falciform seta and few short setae scattered from the base to the level of insertion of the hooked-falciform seta; 4) the distal division of the subapical lobe divided into two arms, the proximal arm with 3 setae, which include a hooked seta (*h*), a saberlike seta (*s*), and a foliform seta (*l*); distal arm with 5 setae, which include a saberlike seta (*s*) and 4 narrow, appressed setae (*f*); 5) the lobes of tergum IX small, cone-shaped, widely separated and with few setae.

In the adults, *Cx. carincii* can be easily separated from *Cx. epanastasis*, *Cx. extenuatus* and *Cx. pedroi* by the hindtarsomeres completely dark-scaled, and from *Cx. adamesi* by the pleural integument with conspicuous pattern of dark spots. *Culex carincii* is morphologically closer to *Cx. crybda* and *Cx. ribeirensis* than to any other species of *Melanoconion*. In the male genitalia, *Cx. carincii* can be separated from all the other species of the Pedroi Subgroup by 1) the distinctive shape of the gonostylus, and 2) the gradual insertion of setae on the distal arm of the distal subapical lobe of the gonocoxite. Furthermore, comparison of the COI sequences of *Cx. carincii* with the ones of *Cx. crybda* and *Cx. ribeirensis* from Brazil [21] showed a mean interspecific divergence of 8.42% and 9.21%, respectively, coherent with three distinct species.

The shape of the gonostylus and IX tergite lobes illustrated for *Cx. crybda* in Sallum and Forattini [26] are very different from the holotype of *Cx. crybda* deposited in the USNM. The distinctive gonostylus and IX tergite lobes contradict the general statement that male genitalia of *Cx. crybda* are identical to the ones of *Cx. adamesi*, *Cx. pedroi*, and *Cx. ribeirensis*. The holotype male genitalia of *Cx. crybda* was never described in detail and the most recent illustrations appear to belong to different species [26; 27]. However, the illustration in Pecor et al. [27] is the one that best matches the holotype male genitalia of *Cx. crybda* deposited in the USNM. The Vomerifer Group was not retrieved as monophyletic in the phylogenetic study of Torres-Gutierrez et al. [36], because two females identified as *Culex* near *gnomatos* and *Culex* near *portesi* were though to belong to this group. Based on our analysis, their *Culex* near *gnomatos* clustered in the same BIN of our male specimens of *Cx. ocossa* of the Ocossa Group, and their *Culex* near *portesi* close to the BIN of our male specimens of *Cx. carincii* of the Crybda Group. This strengthens the hypothesis that these females do not belong to the Vomerifer Group, thus explaining why they clustered outside of it in their phylogenetic analyses.

In the adult, *Cx. extenuatus* can be easily separated from *Cx. adamesi*, *Cx. carincii*, *Cx. crybda* and *Cx. ribeirensis* by the hindtarsomeres with conspicuous white rings at joints. In the male genitalia, *Cx. extenuatus* can be separated from *Cx. epanastasis* and *Cx. pedroi* by the longer and thinner arms of the distal subapical lobe of the gonocoxite. While this diagnostic character seems inconspicuous, we found no intergradation among the material examined. In French Guiana, *Cx. pedroi* was only collected along the coastal plain in swampy ecosystems, while *Cx. epanastasis* and *Cx. extenuatus* were collected along more inland riverine ecosystems where *Cx. pedroi* is apparently absent. Furthermore, molecular delimitation of these three species was corroborated by all the methods, *Cx. extenuatus* showing a mean interspecific distance of 8.13% with *Cx. epanastasis* and 8.14% with *Cx. pedroi*. Comparison of our COI sequences with the ones in Torres-Gutierrez et al. [21] suggests that *Cx. extenuatus* also occur in Brazil. The possibility of a cryptic species among *Cx. pedroi* was already proposed based on the study of the ITS2 ribosomal marker [37]. Unfortunately, our data cannot permit to answer if *Cx. extenuatus* described herein is the *Cx. pedroi*-Peru form of Navarro and Weaver [37]. Species of the Pedroi Subgroup have been recognized as vector of several viruses. For example, *Cx. adamesi* is a natural proven enzootic VEEV vector in Colombia [38], *Cx. crybda* was found naturally infected by Bussuquara and Guama viruses in Panama [39], and *Cx. pedroi* is both a natural proven enzootic VEEV vector in Colombia [38] and the main vector of VEEV in the Peruvian Amazon [40]. The existence of more species than initially thought in the Pedroi Subgroup may explain differences in vector competence between populations with possible outcomes regarding transmission and emergence of pathogens.

### Conclusions

The COI barcode prove again its usefulness and effectiveness in identifying mosquitoes, including cryptic diversity like exemplified here with *Culex* species of the Pedroi Subgroup of the Spissipes Section of *Melanoconion*. However, this study points the limits of this marker for some groups of species within the subgenera *Culex* and *Melanoconion*, probably because of introgression or incomplete lineage sorting. At the scale of South America, inconsistencies between morphological identification and molecular delimitation most likely the result of imperfect taxonomy and geographical gap in sampling. Nevertheless, the COI barcode remains a powerful tool to mitigate morphological taxonomy across geographical scales and efforts should be pursued to improve the delimitation of Neotropical species. Besides adding more taxa, future work should include additional markers to improve delimitation of species poorly supported by the COI barcode alone, and statistically resolve basal nodes toward a more natural classification. This study represents an important contribution to the barcoding initiative of South American mosquitoes and open exiting perspectives to accurately identify *Culex* female and immature specimens during entomological, ecological, or arboviral surveys.

## Acknowledgments

We would like to thank the Office Nationale des Forêts (ONF) and the Parc Amazonien de Guyane (PAG) and more particularly the Réserve Naturelle Nationale de La Trinité, the Réserve Naturelle Nationale des Nouragues (including the Station scientifique des Nouragues), and the Réserve Naturelle Nationale Mont Grand Matoury for logistical support during field studies. We are also thankful to David Pecor from the Walter Reed Biosystematic Unit at the Smithonian Institution for providing images of the holotypes of *Cx. crybda* and *Cx. pedroi*. Finally, we thank the many contributors who helped us to collect some of the voucher specimens during the past 10 years, by first name alphabetical order: Agathe Chavy, Arthur Kocher, Céline Leroy, Clémence Mouza, Estelle Chabanol, Fabienne Jourdan, Gabrielle Georgeon, Guillaume Lacour, Hadrien Lalagüe, Katy Heu, Marceau Minot, Mathilde Gendrin, Ottavia Romoli and Yanouk Epelboin. This study was funded by the “Investissement d’Avenir” grant managed by the Agence nationale de la recherche (ANR) through the “MicroBIOMES” project (CEBA, ref. ANR-10-LABX-25-01), as well as by the Fonds européen de développement régional (FEDER) grant through the “CONTROLE” project (ref. GY0010695, FEDER/2017/N°7).

## Author Contributions

Conceptualization: ST AG BdT AL, RC, PG, JI, ID, JBD. Data Curation: ST AG.

Formal Analysis: ST.

Funding Acquisition: BdT AL ID JBD. Investigation: ST AG BdT RC PG JI JBD. Methodology: ST AG JBD.

Project Administration: ID JBD. Resources: ST JBD.

Software: ST. Supervision: ID JBD. Validation: ST. Visualization: ST.

Writing – Original Draft Preparation: ST AG JBD. Writing – Review & Editing: BdT AL, ID.

## Supporting information captions

**S1 Table. List of the *Culex* species corresponding to the voucher specimens that were COI sequenced in this study.** Species are listed alphabetically by subgenus and the life stage is indicated for each taxa (M: male with dissected genitalia; F: female; L: larva). (DOCX)

**S2 Table. List of Barcode Index Numbers (BINs) with their associated *Culex* species obtained from BOLD (last visited august 2024).** Species are listed alphabetically by subgenus except when more than one morphological species are included in a BIN. Distances (p-distance) correspond to the percentage of dissimilar pairwise nucleotides and counts correspond to the number of voucher specimens included in this study followed, between brackets, by the total number of specimens (including ours) present in the BOLD database. (DOCX)

**S3 Fig. Neighbor Joining tree of 529 COI sequences belonging to 150 species/morphospecies of *Culex* from South America.** This dataset is composed of 246 sequences from French Guiana, 164 sequences from Brazil [17; 21], 62 sequences from Colombia [20], 46 sequences from Argentina [17; 22], and 11 sequences from Ecuador [16]. For each specimen, we indicated the morphological identification, the BOLD specimen code, the original specimen code, the sampling country and the BOLD Barcode Index Number (BIN). Identification of eight specimens from Brazil were missing in BOLD. Identifications of these specimens in the original publication [21] were as follows: KX779777: *Culex akritos*, KX779846: *Culex* nr. *pedroi*, KX779796: *Culex dunni*, and KX779874‒KX779877 and KX779843: *Culex vaxus*.

